# AddBiomechanics: Automating model scaling, inverse kinematics, and inverse dynamics from human motion data through sequential optimization

**DOI:** 10.1101/2023.06.15.545116

**Authors:** Keenon Werling, Nicholas A. Bianco, Michael Raitor, Jon Stingel, Jennifer L. Hicks, Steven H. Collins, Scott L. Delp, C. Karen Liu

## Abstract

Creating large-scale public datasets of human motion biomechanics could unlock data-driven breakthroughs in our understanding of human motion, neuromuscular diseases, and assistive devices. However, the manual effort currently required to process motion capture data and quantify the kinematics and dynamics of movement is costly and limits the collection and sharing of large-scale biomechanical datasets. We present a method, called AddBiomechanics, to automate and standardize the quantification of human movement dynamics from motion capture data. We use linear methods followed by a non-convex bilevel optimization to scale the body segments of a musculoskeletal model, register the locations of optical markers placed on an experimental subject to the markers on a musculoskeletal model, and compute body segment kinematics given trajectories of experimental markers during a motion. We then apply a linear method followed by another non-convex optimization to find body segment masses and fine tune kinematics to minimize residual forces given corresponding trajectories of ground reaction forces. The optimization approach requires approximately 3-5 minutes to determine a subject’s skeleton dimensions and motion kinematics, and less than 30 minutes of computation to also determine dynamically consistent skeleton inertia properties and fine-tuned kinematics and kinetics, compared with about one day of manual work for a human expert. We used AddBiomechanics to automatically reconstruct joint angle and torque trajectories from previously published multi-activity datasets, achieving close correspondence to expert-calculated values, marker root-mean-square errors less than 2 cm, and residual force magnitudes smaller than 2% of peak external force. Finally, we confirmed that AddBiomechanics accurately reproduced joint kinematics and kinetics from synthetic walking data with low marker error and residual loads. We have published the algorithm as an open source cloud service at AddBiomechanics.org, which is available at no cost and asks that users agree to share processed and de-identified data with the community. As of this writing, hundreds of researchers have used the prototype tool to process and share about ten thousand motion files from about one thousand experimental subjects. Reducing the barriers to processing and sharing high-quality human motion biomechanics data will enable more people to use state-of-the-art biomechanical analysis, do so at lower cost, and share larger and more accurate datasets.

## Introduction

Quantitative analysis of human movement dynamics is a powerful tool that has been widely used to estimate joint loading during walking and running e.g. [1–9], assess muscle function during gait in individuals with cerebral palsy e.g. [10, 11], analyze the performance of assistive devices for improving human movement e.g. [12–15], quantify changes in neuromuscular control due to Parkinson’s disease e.g. [16, 17], and even generate more realistic computer graphics e.g. [18–20]. But the resource-intensive nature of quantitative movement analysis restricts access to this data and keeps study sample sizes small. Without automated tools to process, analyze, and harmonize lab-based human movement data, the biomechanics field has been hamstrung in its ability to apply modern, data-hungry machine learning approaches to create accurate, data-driven models to predict, prevent, and personalize treatment for the many injuries and conditions that impair movement.

Laboratory-based motion capture is the current benchmark data acquisition technique to quantify human biomechanics [21, 22], but current state-of-the-art software for reconstructing the motion and kinetics of a human musculoskeletal model from optical marker trajectories and ground reaction forces requires substantial iterative “guess-and-check” refinement, which increases costs, limits scalability, and reduces the reproducibility of motion capture studies [23–25]. A typical experiment involves placing optical markers on a subject’s body segments and having the subject perform actions in a laboratory space surrounded by specialized cameras. These camera systems and associated software are able to reconstruct the three-dimensional locations of the optical markers in the lab, and given the marker trajectories over time, one can use proprietary, open, or custom software to reconstruct the kinematics of the subject’s body segments. If external loads recorded simultaneously from ground force plates an inverse dynamics method can be used to estimate the joint torques the subject used to generate the observed motion.

### Current practices for model scaling and inverse kinematics

To reconstruct movement kinematics from optical motion capture data, software must address several sources of noise, ambiguity, and model error. Given a set of marker trajectories corresponding to a motion of interest, software must reconstruct a digital twin of the experimental subject, with segment dimensions that match the subject as closely as possible. This process is called *scaling*, and a variety of approaches have been described [23, 26–33]. Finding accurate scaling is especially important when using motion capture data to create muscle-driven simulations because the muscle-tendon parameters are scaled by the body segment dimensions [34]. To achieve accurate kinematic results, the locations of the markers on the scaled digital twin must be adjusted to account for variations caused by human error in attaching the markers to the body and the variations in the dimensions of human subjects [24]. This is called *marker registration*. Finally, the positions and orientations of the body segments over time must be determined, which is typically done using an optimization process called inverse kinematics [35–39]. Inverse kinematics algorithms generally produce more accurate results when the solutions are constrained by an underlying skeletal model [13, 24, 40].

The interdependence between scaling, marker registration, and inverse kinematics means that experts must follow an iterative guess-and-check procedure, where they refine each of the steps several times, making small adjustments to each value until a desired accuracy is achieved [41, 42]. For example, increasing the length of the upper arm segment in a subject’s digital twin will require also changing the marker registrations for any markers on the forearm and the hands, because otherwise those markers would move as a result of the longer upper arm. A longer upper arm will also, all else being equal, change the resulting motion found by inverse kinematics. While there are best practices for conducting validation at each step [34], the process typically requires extensive and subjective input from an expert.

Automating the scaling and registration process has been studied before, in pioneering work by Reinbolt et. al. [43] and Charlton et. al. [44]. These authors used gradient-free optimization methods to automatically estimate body segment scales and marker registrations while solving gradient-based inverse-kinematics problems repeatedly in an inner-loop to evaluate optimization progress. These methods require large amounts of compute time because every iterative guess the outer optimizer makes about body segment scaling and marker offsets requires solving a computationally costly inner optimization problem (inverse kinematics) to evaluate the quality of the guess. The method of Reinbolt et al. [43] produces the best results using a particle-based optimizer for their outer optimization problem, to combat the non-convexity of the problem, but this comes at a further increase in computational cost.

Given the interconnected nature of body segment scaling, marker registration, and inverse kinematics, one might also consider posing all three problems as a single optimization problem. However, such a formulation leads to a nonconvex optimization in which a global solution is not guaranteed [45]. Instead, we can only guarantee to find a local optimum close to an initial guess, so providing a high quality initial guess is crucial. Andersen et al. [46] have formulated such nonconvex optimization problems, but did not address the problem of reliably finding an initial guess for the non-convex optimization problem proposed.

Markerless motion capture systems based on video recordings have recently become popular since they do not require expensive motion capture equipment [47]. While these approaches do not track optical markers, recent work has focused on combining markerless motion capture techniques (e.g., pose detection) with scaled musculoskeletal models to incorporate physiological joint constraints [48, 49]. These approaches still rely on solving an inverse kinematics problem, using keypoints from pose detection algorithms, rather than optical markers. Accurate scaled models also enable deeper biomechanical analyses with markerless motion capture techniques to estimate kinetic quantities, like joint moments and muscle forces [49].

### Creating physically-consistent simulations

Making accurate conclusions about the kinetics of human movement requires that the kinematics and mass properties of a musculoskeletal model are “dynamically-consistent” with external forces (e.g., ground reaction forces). Incorporating experimental, external force measurements into simulations of movement can lead to challenges similar to those presented in the scaling and inverse kinematics problems. When inconsistencies between model properties, kinematics, and measured external forces are present, an inverse dynamics analysis will yield physically impossible external forces and moments about the model’s root segment (e.g., pelvis), often referred to as *residual forces*. Biomechanics researchers aim to minimize or eliminate residual forces and moments from their simulations; in practice, it is usually sufficient to reduce the magnitude of the residual forces below recommended thresholds based on the magnitude of the experimental ground reaction forces and center of mass trajectory [34].

Similar to model scaling, dynamic consistency is usually achieved through an iterative process where changes in model kinematics and mass parameters are made to reduce residual forces and moments. OpenSim, widely used simulation software, provides the Residual Reduction Algorithm (RRA) tool, which adjusts mass, body mass center locations, and joint kinematics to minimize residual forces and moments [41, 50]. The RRA tool uses a tracking controller to adjust joint kinematics while penalizing the magnitude of residual forces and moments. Tracking weights for each joint must be chosen such that the kinematic changes are within measurement errors while still minimizing residual forces. Since changes in residual forces are dependent on changes in kinematics and mass properties, it is often necessary to run the RRA tool iteratively to meet recommended residual force thresholds. Sturdy et al. [25] used the RRA tool to automate the reduction of residual forces by optimizing the tracking weights with random hill climbing. This approach yielded residuals within recommended thresholds from Hicks et al. [34], but required a pre-scaled model, joint trajectories from inverse kinematics, and up to 2 hours of processing time per subject on a standard desktop machine.

### Automating motion capture data processing with AddBiomechanics

Thus, despite recent advances in biomechanics simulation methods, reconstructing human movement from experiments remains a challenging and time-consuming task for researchers, and large-scale datasets are lacking. This paper introduces an automated method (Fig 1), called AddBiomechanics, that uses a combination of traditional kinematic solvers and modern bilevel optimization to estimate high quality inverse kinematics and dynamics from experimental motion capture data in reasonable computation time. We first apply a sequence of optimizations to approximate the initial values for each of the body segment scales, marker registrations, and inverse kinematics [43, 44]; thus, no user-provided initial guess is required. Then, rather than iteratively repeat those optimization problems hundreds of times as in previous work, we apply bilevel optimization techniques to simultaneously optimize body scaling, marker registration, and inverse kinematics. Next, we find a least-squares fit for the subject mass and initial center-of-mass position and velocity such that integrating the center-of-mass accelerations (which are the measured ground-reaction-forces divided by subject mass) results in the least-squares closest approximation to the purely kinematic motion we found in the previous step. Finally, we optimize body segment masses and tune the body scales, marker registrations, and model kinematics using the same bilevel approach to find a motion that is still consistent with the experimental marker data while achieving nearly zero residuals. To evaluate the algorithm, we computed marker RMS errors and residual forces and moments for a set of common movements studied in the biomechanics field including walking, running, squatting, and sit-to-stand motions, and compared errors to results computed by experts. We also used AddBiomechanics to estimate joint angles and moments for a simulated walking motion with known dynamics and zero residuals. Finally, we evaluated the computational cost of computing kinematics and kinetics on these datasets.

**Fig 1.**
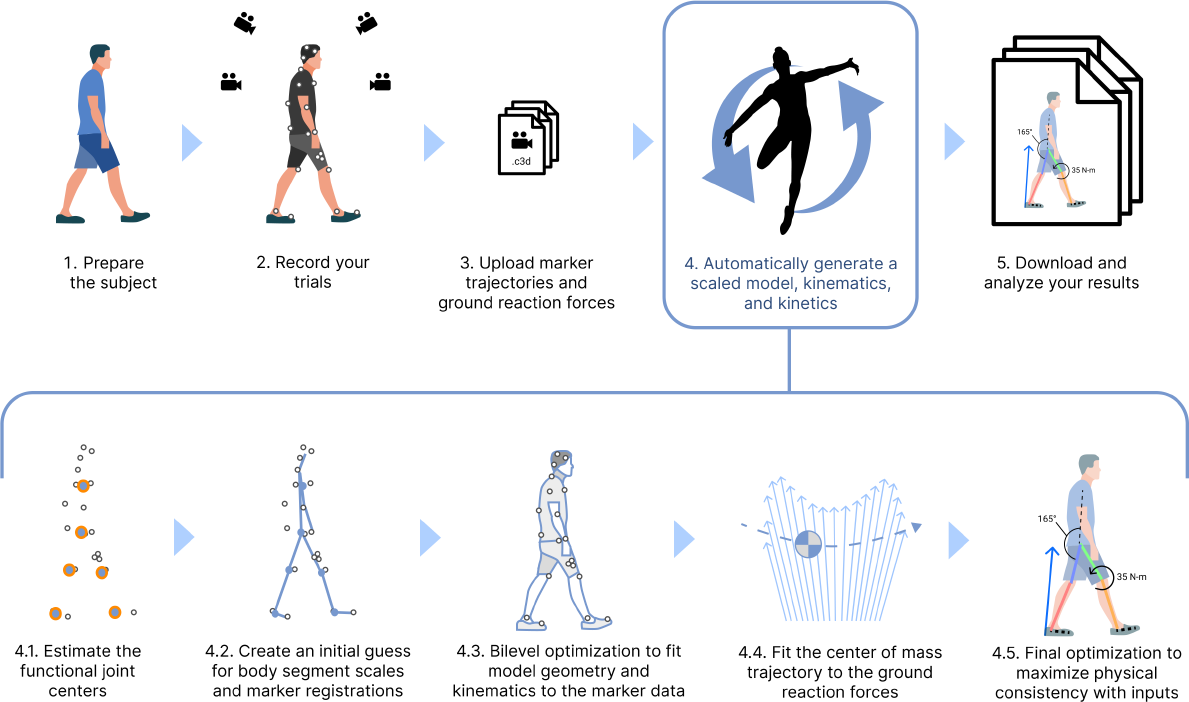
AddBiomechanics automates the analyses required in a standard motion capture pipeline. AddBiomechanics integrates into the standard motion capture pipeline to automate the process of model scaling, marker registration, inverse kinematics, and residual reduction. Once experimental marker and ground reaction force data have been collected and uploaded (steps 1-3), AddBiomechanics (step 4), replaces time-consuming and error-prone manual steps in previous workflows. Our method processes input marker and force data through several steps automatically. First, it finds the functional joint centers from the data (step 4.1), and then it uses the marker data and those joint centers to make an initial guess for body segment scales and marker registrations (step 4.2). The initial guess then serves as the starting point for a bilevel optimization problem that matches the model geometry and kinematics to the experimental marker data as closely as possible (step 4.3). Next, the model trajectory is updated by fitting the center of mass motion to the ground reaction force data (step 4.4). A final optimization adjusts body segment masses and joint kinematics to maximize consistency between the model and the experimental data (step 4.5). The final output is a musculoskeletal model scaled to the subject with registered markers, joint angles, and joint torques over time.

AddBiomechanics can process large amounts of motion capture data automatically. To facilitate its use, we have released the software as an open source cloud-based service available at AddBiomechanics.org, where over 300 researchers from dozens of institutions have begun to process their data without downloading or installing any software. AddBiomechanics outputs OpenSim project files [41], compatible with the widely used open source biomechanics package, so the results of scaling and marker registration can be transferred to OpenSim for further analysis. Optimized skeletons can also be exported in formats compatible with MuJuCo [51] and PyBullet [52], which are physics simulators commonly used in reinforcement learning and computer graphics.

## Methods

Given a musculoskeletal model and experimental data, AddBiomechanics solves a sequence of optimization problems to compute model scaling, inverse kinematics, and inverse dynamics, where the solution for each problem is the initial guess for the subsequent problem. First, the model scaling and inverse kinematics problems are solved using a series of linear and bilevel optimization problems to find a solution for the model body segment scale factors, marker registrations, and joint kinematics. If ground reaction force data is provided by the user, AddBiomechanics then estimates center of mass trajectory and overall subject mass with a linear optimization, followed by a non-convex optimization step to minimize residual forces and tune the original model scaling and joint kinematics solution. Each of these steps are described in more detail in the sections that follow.

### Input model and experimental data

#### Generic, unscaled musculoskeletal model

Our algorithm can scale and register markers on arbitrary skeletons defined using the OpenSim model format. A skeleton is composed of a set of body segments, connected by joints. The scaling of each link is concatenated to form the ***s*** vector, and the degrees of freedom of each joint are concatenated to form the ***q*** vector. The algorithm supports all OpenSim joint types, including custom joints. Examples of skeletons that have been successfully scaled and registered in our experiments include widely used state-of-the-art biomechanical models [53, 54].

#### Motion capture marker trajectories

The output of a commercial motion capture system is a series of frames, often at 100-200 Hz, where each frame contains 3D coordinates representing the trajectories of optical motion capture markers in the experimental capture volume at the corresponding moment in time. Users must provide these marker trajectories for each experimental movement trial. Each 3D coordinate must be “labeled” with a tag corresponding to an experimental marker location on the subject (e.g. “C7” for the optical marker placed on the C7 spinal segment). A full list of marker tags, and their location on a given musculoskeletal model is known as the “marker set.” We provide models with default marker sets, but users may upload a custom model with a marker set that matches the experimental marker data they provide. In practice, markers are almost never placed exactly at their ideal locations, and these small deviations in experimental marker placement must be accounted for during the marker registration step. Not every marker from the marker set is observed in every frame, because markers may be occasionally obstructed during a motion capture experiment. Our algorithm allows for markers with missing frames and can automatically adjust for deviations in marker placement during the optimization.

#### Ground reaction forces

Ground reaction forces are recorded from force plates embedded in the ground and are typically measured at higher frame rates (e.g., 1000-2000 Hz) compared to marker trajectory measurements. To compute dynamics with AddBiomechanics, users must provide the 3 forces, 3 torques, and center of pressure locations for each force plate as a C3D file or tab-delimited data file. We assign loads from each force plate to the feet in the model based on when the feet are penetrating the ground within known force plate geometries and when the ground reaction force information exceeds a non-zero threshold. We assume that both feet are never simultaneously in contact with a single force plate.

## Model scaling and inverse kinematics

### Model optimization to minimize marker position errors

Given the measured marker trajectories from a motion capture system with length equal to the number of time points *T*, 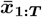, AddBiomechanics formulates a nonconvex optimization that solves for the kinematic pose trajectories, ***q***_1:*T*_, the scaling parameters of the body segments of the musculoskeletal model, ***s***, and the locations of markers attached to the body segments, ***p***. The objective of the optimization is to minimize the deviation of estimated marker positions from 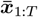:

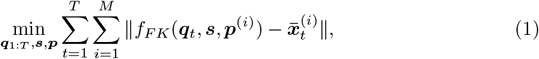

where *M* is the number of markers, ***p***^(*i*)^ *∈* ℝ^3^ denotes the position of the *i*-th marker in the local frame of the body segment to which it is attached, and ***p*** *∈* ℝ^3*×M*^ is the concatenated local positions of all markers. *f*_*F K*_(***q***_*t*_, ***s, p***^(*i*)^) is the forward kinematic process that transforms a point ***p***^(*i*)^ in a skeleton scaled by ***s*** and in the pose ***q***_*t*_ from the local coordinate frame of the assigned body segment to the world coordinate frame. Note that we use the *t* to denote the time index (rather than a value in seconds) throughout the manuscript.

Eq (1) is high-dimensional and nonconvex. Consequently, the solution of such an optimization is highly sensitive to the initialization of the decision variables. We use a bilevel maximum-a-posteriori (MAP) optimization and an initialization strategy to achieve new state-of-the-art in automatic processing of biomechanical motion capture data. The proposed bilevel MAP optimization simultaneously considers data reconstruction and anthropometric statistics when jointly optimizing all decision variables in Eq (1). To overcome the sensitivity to the initial guess, our method individually initializes each type of variable using independent sources of information. Specifically, we use kinematic constraints to initialize ***q***_1:*T*_, a geometric invariant to initialize ***s***, and real-world measurement to initialize ***p***. Once the variables are initialized individually, the final bilevel optimization ensures that they agree with one another, given the observed data and model priors. More details about each of these steps are provided in the sections that follow.

### Bilevel maximum-a-posteriori (MAP) optimization

Given recorded marker positions 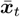 at time index *t*, we are interested in reconstructing the scales of each body segment in the musculoskeletal model, ***s***, the local positions of the markers ***p*** attached to their assigned body segments, as well as the joint pose ***q***_*t*_. This problem can be formulated as a maximum a-priori (MAP) optimization:

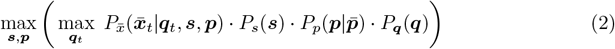

The first term, 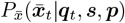, is a conditional probability of the observed data given the estimated parameters. This formulation is equivalent to the standard least-squares inverse kinematics objective term if we assume Gaussian noise in our marker observations. The second term, *P*_*s*_(***s***), expresses the prior of skeleton scaling, encoded as a multivariate Gaussian fit to the ANSUR II dataset [55] of anthropometric scalings. If the height, weight, or biological sex of the experimental subject is known, the multivariate Gaussian skeleton scaling prior is conditioned on that information before any optimization. The third term, 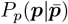, is a zero-mean Gaussian distribution that regularizes the deviation of the marker locations from their intended locations 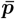 provided by the experimenter, encoding that markers are generally placed close to their intended locations, even if they do not perfectly align. 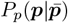 regularizes markers differently: some markers are placed on anatomical landmarks, and therefore are unlikely to move relative to the landmark from subject to subject, and other markers are placed anywhere on a body segment as “tracking” markers, and therefore the optimizer should be allowed wide discretion to adjust those marker locations. The sets of “anatomical” and “tracking” markers are determined from the musculoskeletal model provided by the user. For best performance, users should place at least one anatomical marker on each body segment in the model. The fourth term is a prior over ***q***, but we assume this is a uniform distribution and drop it hereafter.

This is a bilevel optimization problem, because in order to evaluate the quality of given skeleton scaling ***s*** and marker locations ***p***, we need to optimize over the possible joint positions ***q***_*t*_. To efficiently solve the bilevel optimization problem, we observe that at the optimal values of ***q***_*t*_ for max_***q****t*_ 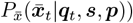, the gradient of the inner optimization problem will be zero. Using this observation, we reformulate the bilevel optimization problem as a single-level nonconvex optimization problem with nonconvex constraints. For numerical stability, we minimize the negative log of the above objective function:

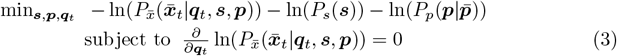

At a locally-optimal point, the gradient of the objective term with respect to any of the decision variables is zero, so it must be zero with respect to ***q***_*t*_:

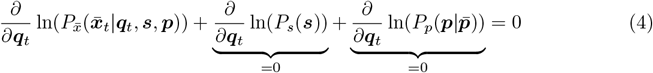

Thus, at a locally-optimal point for the objective function, the constraint in Equation 3 must hold regardless, and so we could theoretically omit it from the optimization problem without loss of correctness. However, we found that explicitly including the constraint allows the optimizer to converge to a high-quality solution much more quickly. See S1 Appendix for a more detailed analysis.

We could use any nonlinear optimization solver to solve Eq (3). In practice, we use IPOPT [56], which is a high-quality and open source solver. However, due to our problem’s non-convexity, a good initial guess for the decision variables is needed to produce reasonable results.

### Initializing the kinematic decision variables

Prior to solving the optimization problem in Eq (3), we need to get “close-enough” initial guesses for the decision variables. We do this through a sequence of optimization problems as described in the steps below. We obtain initial guesses for the joint angles, ***q***_*t*_, body segment scales, ***s***, and marker offsets, ***p***, individually based on independent sources of information such that the cascading errors can be mitigated.

1. Initialize ***p*** using the marker locations measured by the experimenter or defined by the existing marker set.
2. Initialize ***s*** by analytically computing the functional joint centers and axes using the method described in [57], refine those values using a non-convex sphere-fitting problem, and scale ***s*** to match the joint axes along with the measured markers.
3. Initialize ***q***_*t*_ by solving inverse kinematics with a skeleton scaled to ***s*** and with marker locations ***p***.

Step 1 is trivial and Step 3 is a simplified Eq (1) with ***s*** being given from Step 2 and ***p*** given from Step 1, min_***q****t*_ 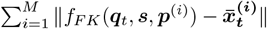. Solving this inverse kinematics problem efficiently has been an area of research for decades [58–60] and can be done efficiently and reliably.

The most involved step in our initialization process is Step 2, initializing the body segment scales ***s***. We begin by analytically computing a set of functional joint centers and axes from the measured marker trajectories using the least-squares method given in [57]. The least-squares method is deterministic, but can be slightly less than optimal in the presence of soft-tissue artifacts, so we further refine joint center estimates with a non-convex problem, initialized with the answers we get from [57]. Let the subset of markers attached to the two body segments connected by the joint be ℳ. We can estimate the joint position ***c*** in the world frame over time by

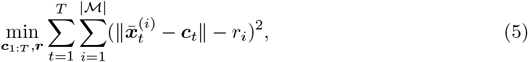

where *r*_*i*_ is the estimated distance between the *i*-th marker 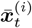 and the joint center ***c***_*t*_ for all *t. r*_*i*_ is constant over time. For each marker, Eq (5) fits a moving sphere centered at ***c***_*t*_ with the radius *r*_*i*_, to match the measured positions of the marker over time.

The sphere-fitting approach to finding functional joint centers can yield ambiguities when marker motion adjacent to a joint is primarily confined to the sagittal plane, as commonly happens in locomotion. In such cases, we could move our joint center perpendicular to the sagittal plane, and still have an equally good solution for sphere-fitting. As a result, sphere-fitting might incorrectly scale the skeleton to match erroneous joint positions. For example, we might incorrectly scale the hip width while still matching all the measured marker motion for the thighs and the pelvis.

To address these ambiguities, we formulate another optimization problem to simultaneously find the joint axis and the joint center, building on Eq (5). This problem is similar in spirit to the axis-of-rotation problem described in [61], but can be implemented without any matrix factorizations. The goal of the axis fit problem is to identify not only a joint center ***c***, but also the direction of axis ***a*** at each frame. We also estimate a fixed distance from the center for each marker, parameterized by a distance *u*_*i*_ along the axis ***a*** and a distance *v*_*i*_ perpendicular to the axis ***a***. The result of a successful axis fit is that we capture a line at each frame, where the functional joint center could lie anywhere on that line:

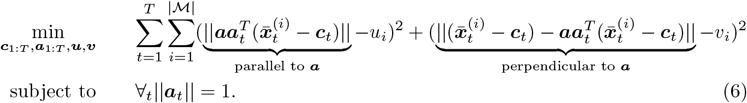

For each marker 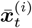 in the set *ℳ*, we decompose 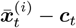 to two vectors: the parallel vector which is the projection of 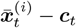 on ***a***_*t*_, and the remaining orthogonal vector. Eq (6) encourages that both the projected vector and the orthogonal vector maintain constant length over time for every marker in ℳ.

We run both sphere fitting and axis fitting at each joint. Because the axis fit is a strictly more demanding problem, if it succeeds, then the axis is passed on as a constraint for subsequent problems. If axis fitting fails, then it must be because there is out-of-plane marker motion, which means that the sphere fit is not ambiguous, so then the exact joint center is passed along to subsequent problems.

Once we determine the joint center and/or the joint axis, we formulate another optimization to initialize the scaling parameters ***s***:

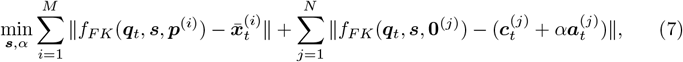

where the zero vector **0**^(*j*)^ indicates the local coordinate of the joint *j*, and *N* is the number of joints. The first term fits the skeleton to the measured marker positions, while the second term encourages the joints to lie on the estimated joint axes solved by Eq (6), at a distance controlled by the scalar decision variable *α*. If the joint axis does not exist for the joint *j*, we set ***a***_*j*_ to zero and remove *α* from the optimization.

After initializing the decision variables, we find a solution by minimizing Eq (3). The body scales, ***s***, and marker registrations, ***p***, are returned to the user as an optimized version of the OpenSim model the user submitted to the tool. The joint angle trajectories, ***q***_*t*_, obtained from inverse kinematics solution for each trial are exported using OpenSim’s MOT file format.

## Inverse dynamics

### Model optimization to achieve physical consistency

After finding a set of marker registrations and body scales that achieve a good inverse kinematics fit to the marker trajectories, we can then solve another optimization problem to find body segement masses and updated joint kinematics that minimize the set of residual forces and torques applied to the pelvis. Similar to the model scaling and inverse kinematics optimization, this problem is non-convex, so we first need to create a good initial guess for the model and mass parameters and update the kinematic trajectory so that it is physically consistent with the observed ground reaction force data. To achieve this, we solve a series of linear equations to fit the system’s center of mass trajectory to our results from the marker fitting step while prescribing the observed ground reaction forces. We begin with the solution ***q***_1:*T*_ obtained from the previous model scaling and inverse kinematics process. To avoid large acceleration artifacts in the dynamic fitting problems, we smooth the solution ***q***_1:*T*_ by minimizing the jerk of the joint angle trajectories over time (S1 Appendix).

### Center of mass trajectory fitting

The trajectory of the center of mass of the system is dictated by the ground reaction forces acting on the model and can be defined by the differential equation:

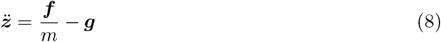

where 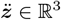 is center of mass acceleration, ***f*** is the ground reaction force vector, *m* is the system mass, and ***g*** is gravitational acceleration.

Since the ground reaction forces are known from experimental data, the center of mass acceleration is just a linear function of *inverse* mass of the model. We define a new variable 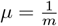, and note that the center of mass trajectory is a linear function of *μ*. If the initial state (the state at index *t* = 1) of the center of mass acceleration, (***z***_1_, ***ż***_1_), is known, the entire trajectory ***z***_*t*_ is determined. We aim to find a best fit of this trajectory to the trajectory that we obtained from the marker-fitting optimization, 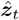.

We define a vector ***ζ*** that contains the three unknown quantities:

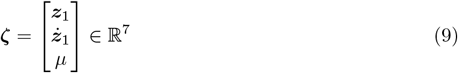

We can define a linear system with matrix ***A*** *∈* ℝ^3*T ×*7^ and offset ***b*** *∈* ℝ^3*T*^ that maps the vector ***ζ*** onto 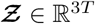, a vector of concatenated center of mass position vectors over time:

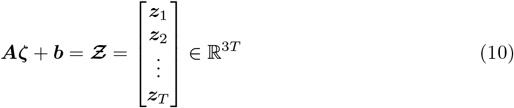

Given the observed trajectory of center of mass motion from the marker fitting step, 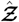, it is possible to find a least-squares best estimate for the unknowns, 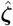, using the pseudo-inverse of ***A***:

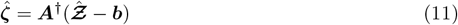

To derive ***A***, first, we define a semi-explicit Euler integration scheme to solve for the center of mass trajectory:

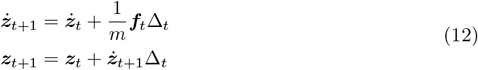

where Δ _*t*_ is the integration time step in seconds.

We can then construct ***A*** and ***b*** using this integration scheme to relate the unknowns ***ζ*** to the center of mass positions, 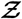:

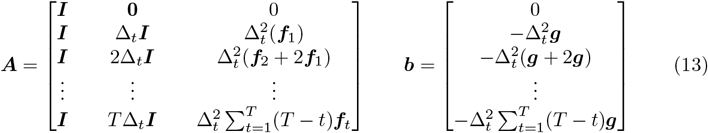

Here, the first two columnar blocks of ***A*** represent the contributions from ***z***_1_ and ***ż*** _1_ to the trajectory 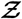, where ***I*** and **0** are the 3 *×* 3 identity and zero matrices, respectively. The third columnar block of ***A*** represents the contribution from the inverse mass *μ* and is a single column containing terms corresponding to the time integration of the ground reaction forces. Similarly, the vector ***b*** contains terms corresponding to the time integration of gravitational acceleration.

By solving Eq (11), we obtain a least-squares best fit of the initial conditions and mass of the system, 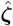, and can use this solution to obtain a new trajectory for the center of mass that is physically consistent with the observed ground reaction force data, 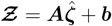. We can recover total mass as 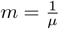. Finally, we modify the position of the pelvis over time while keeping the remaining joint angles fixed to update the model’s center of mass trajectory to match 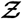. This step serves as an initialization for the final problem described later, which will further refine the joint angle trajectories while optimizing the mass properties of the model.

### Angular dynamics fitting

Fitting the center of mass trajectory provides better physical consistency with the linear ground reaction forces applied to the system, but the trajectory may still be inconsistent with the moments these forces produce about the center of mass of the system. Given our solution to the linear center of mass fitting problem, 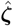, we can expand our approach to also address physical inconsistencies in the angular dynamics.

We use ***θ***_*t*_ *∈* ℝ^3^ to denote the rotational generalized coordinates of the root segment (e.g., the pelvis) at time *t*, which are a subset of the coordinates in ***q***_*t*_. First, we assume that changing ***θ***_*t*_ does *not* change the mass matrix or the Coriolis forces for the skeleton at time *t*. This is not true in general, but since we aim to make small adjustments to ***θ***_*t*_ from the inverse kinematics solution, we find this in practice to be a reasonable approximation when creating an initial guess for the skeleton’s root trajectory. We can then construct a new linear map that relates the initial conditions of the root segment to the trajectory, **Ξ** *∈* ℝ^6*T*^, which includes both the pelvis coordinate rotations, **Θ** *∈* ℝ^3*T*^, and center of mass positions, 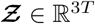 :

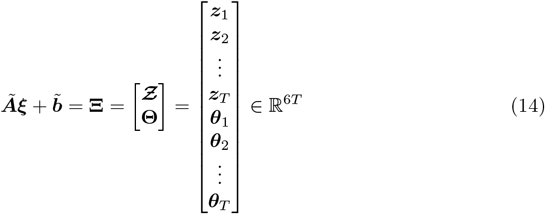

The vector ***ξ*** contains the initial conditions of the center of mass trajectory and the initial pelvis rotational coordinate values, ***θ***_1_ and speeds, 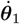:

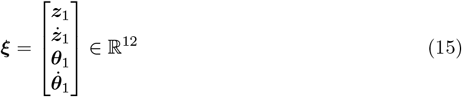

The initial values of ***z***_1_ and ***ż***_1_ are chosen based on our previous solution to the center of mass trajectory fitting problem. Note that unlike in the previous linear fitting problem, we now hold the skeleton mass fixed, so no inverse mass term appears in ***ξ***, and what used to be the third columnar block in ***A*** in Equation 13 is now instead part of the constant term and appears in 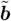. See S1 Appendix for details on how 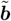 is constructed. As before, we construct ***Ã*** to map the initial conditions ***ξ*** onto the trajectory **Ξ**:

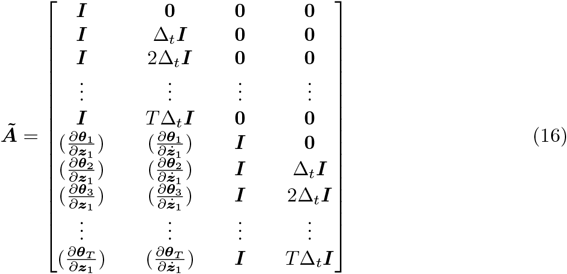

Note that the upper left and lower right quadrants of ***Ã*** are identical to the block matrices we constructed in ***A***, since we use the same semi-explicit integration scheme for both ***z***_*t*_ and ***θ***_*t*_ as defined in Eq (12). The center of mass trajectory ***z***_*t*_ does not depend on ***θ***_1_ or 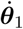, so the upper right quadrant contains all zeros.

To compute the terms 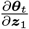 and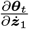 in the lower left quadrant of ***Ã***, we first note that the center of pressure locations are fixed based on the ground reaction force data. Therefore, if we change the location of the center of mass by some finite value Δ***z***_*t*_, the moment applied by the ground reaction force about the pelvis changes by Δ***τ***_*t*_ = Δ***z***_*t*_ *×* ***f***_*t*_. This means that the acceleration of the pelvis rotational coordinates changes by 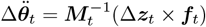, where ***M***_*t*_ is the generalized mass matrix for our skeleton in configuration ***q***_*t*_ found by the inverse kinematics and scaling steps. This can be rewritten as a linear expression between Δ***z***_*t*_ and 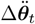 using the skew-symmetric matrix [***f***_*t*_]:

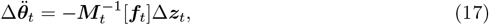

where 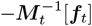 is a constant matrix in ℝ^3*×*3^.

Note that Eq (17) is true for the initial time step even without our simplifying approximations (that changing ***θ***_*t*_ does not effect mass matrix ***M***_*t*_ or Coriolis forces). These approximations are only necessary when we begin to integrate this expression forward in time, since changes in ***θ***_*t*_ will change the mass matrix, ***M***_*t*_, and the linear offsets from the equations of motion (e.g., the Coriolis forces) contained in 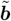, which would render the problem non-linear.

We can now compute 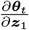 and 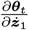 by multiplying together known terms based on the chain rule:

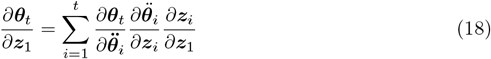

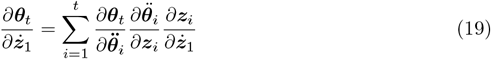

Where the partials are given by:

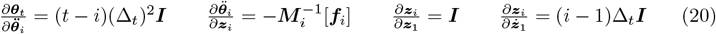

In Eq (19), the first two terms are the same as Eq (18), and the third term is the change in center of mass position due to the change in ***ż***_1_. Both 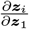 and 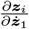 can be obtained directly from ***A***.

The vector 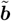 includes terms for the time integration of gravitational acceleration, the acceleration due to the applied ground reaction forces, and the Coriolis terms of the equations of motion of the skeleton. In general, the Coriolis terms depend on ***θ***_*t*_, but based on our simplifying assumption to keep the problem linear, we simply use the initial guess for ***θ***_*t*_ to compute the terms in 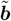. Refer to S1 Appendix for more details on the construction of 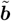.

We can then find a least-squares best fit for the unknown initial conditions, 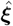, given the observed trajectories of the center of mass position and pelvis rotation coordinates, 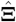, using the pseudo-inverse of ***Ã***:

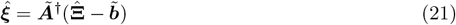

We use the solution 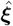 to reconstruct a physically-consistent trajectory for the pelvis coordinate rotations and center of mass positions, 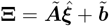. To make the problem linear, we have assumed that our solution for the pelvis coordinate rotations, **Θ**, does not change the mass matrix or Coriolis terms, but since this is not true in general, the solution to Eq (21) will change the terms in ***Ã*** and 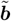 Therefore, to find a satisfactory initial guess for the skeleton’s root trajectory, we form and solve the system defined by ***Ã*** and 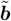 iteratively until **Ξ** converges. In practice, we find that convergence typically takes less than 30 iterations with each iteration taking less than a second on a low-end server.

Once the solution 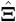 has met our convergence criteria, we have found a trajectory for the center of mass translation and the pelvis coordinate rotations that is physically consistent with the measured ground reaction force data. Finally, we include additional terms to account for errors in force plate locations and orientations and to eliminate drift in very long trials; the details of these terms can be found in S1 Appendix.

### Final optimization to tune marker fitting results and minimize residual loads

After fitting the center of mass trajectory and pelvis coordinate rotations to achieve physical consistency with the ground reaction force data, we run a final optimization to tune skeleton segment masses, marker offsets, segment scale factors, and joint coordinates to minimize the residual forces at the pelvis, 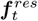, while still retaining a good kinematic fit to the marker data. We achieve this by taking the marker fitting problem described in Eq (2) and adding the segment masses to the decision variables and a loss term to penalize the residual forces:

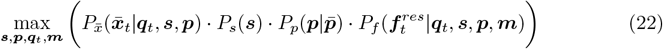

We optimize this problem in the same way as the marker fitting problem, where we minimize the negative log of the objective in Eq (22). Note that we do not use a bilevel problem formulation here, since we now allow the solution to deviate slightly from a valid inverse kinematics solution in order to achieve dynamic consistency. Therefore, we no longer explicitly constrain that the gradient of the inverse kinematics loss term with respect to the joint coordinates be zero.

### Open source implementation

To facilitate adoption, we provide the algorithm as an open-source, cloud-based tool that allows researchers to automate scaling, marker registration, inverse kinematics, residual reduction, and inverse dynamics for their motion capture data without downloading or installing any software, available at AddBiomechanics.org. Users can drag and drop files for automated processing, and then visualize on the web or download results for analysis in OpenSim (Figure 2). C3D or TRC marker files are supported, and C3D or MOT files for ground reaction forces. The cloud tool also allows researchers to automatically generate comparisons of their own hand-scaled data versus the output of the automated system.

**Fig 2.**
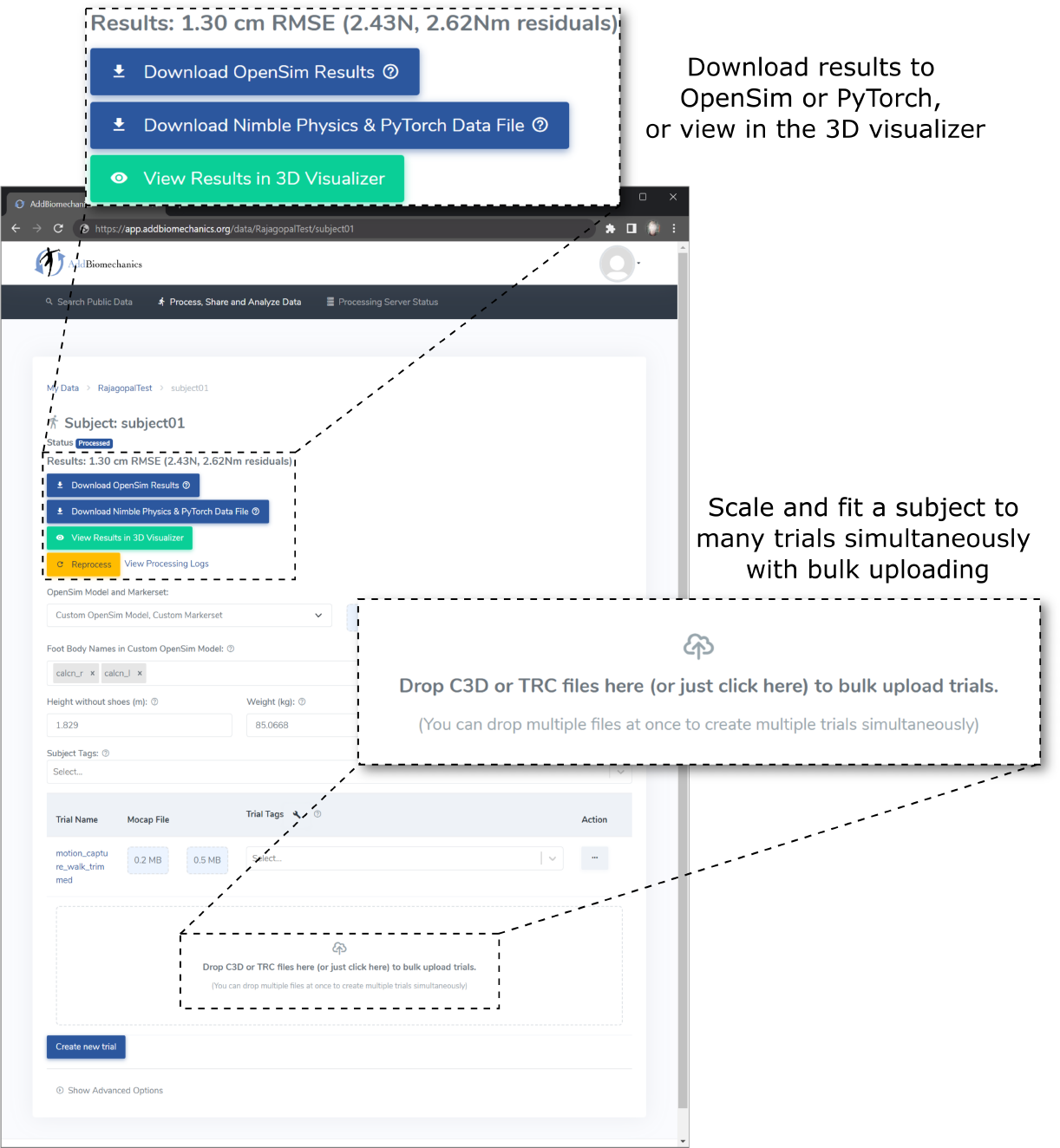
The web interface for AddBiomechanics. The web interface allows users to drag and drop data files for individual experimental trials and the subject data is processed automatically in the cloud.

### Evaluation

To evaluate our algorithm, we first compared AddBiomechanics to expert-computed values for a dataset published by Hamner et al. (2013) with ten subjects running at 2.0, 3.0, 4.0, and 5.0 m s^-1^ [62] (40 total trials), as well as a multi-activity dataset [49] that included sit-to-stand, squatting, jumping, and walking motions (104 total trials). We compared root mean squared errors between experimental and model markers and computed residual forces and moments for both the expert- and AddBiomechanics-determined values. We also qualitatively compared joint angles and joint torques. We used the model, marker set, and raw experimental data (markers and ground reaction forces) from the original study as inputs to AddBiomechanics and compared to the published results computed by the study investigators.

Quantitative comparison of the solved joint angles and moments with ground truth values is another critical test of our method. However, ground truth joint angles and moments cannot be directly measured from experiments. We thus used a three-dimensional dynamic simulation of walking created using trajectory optimization [63], where joint angles and moments are known and residual forces and moments are also known to be zero, to generate a synthetic dataset. We used synthesized marker trajectories, along with the computed ground reaction forces and centers of pressure from the simulation, as inputs to AddBiomechanics. Additional inputs included the original generic, unscaled model and an unregistered version of the appropriate marker set. We then used AddBiomechanics to optimize and compared the recovered motion to the known joint angles and moments.

## Results

### Human expert versus automated processing: running dataset

The average marker RMSE achieved by AddBiomechanics for the running dataset was 1.5 cm, which is significantly smaller than the 4.3 cm marker RMSE (p *<* 0.005, paired t-test) in the originally published results from [62] obtained after using OpenSim’s Residual Reduction Algorithm (Fig 3, left) to modify the running kinematics to reduce residual loads. In addition, the maximum marker error produced by AddBiomechanics (3.8 cm) was smaller than the maximum marker error in the expert-processed results (7.5 cm). AddBiomechanics produced a small but significant reduction in average RMS residual force magnitude (p *<* 0.05, paired t-test) compared to the original study (Fig 3, right). In addition, AddBiomechanics was able to significantly reduce residual torque magnitudes (p *<* 0.005, paired t-test) such that they were below the threshold recommended by Hicks et al. [34], which was not achieved in the original study. Finally, the lower-limb joint angle and joint torque trajectories from the automated approach were qualitatively similar to the trajectories from the original study (Fig 4). AddBiomechanics produced similar results in both the stance and flight phases of running across all subjects.

**Fig 3.**
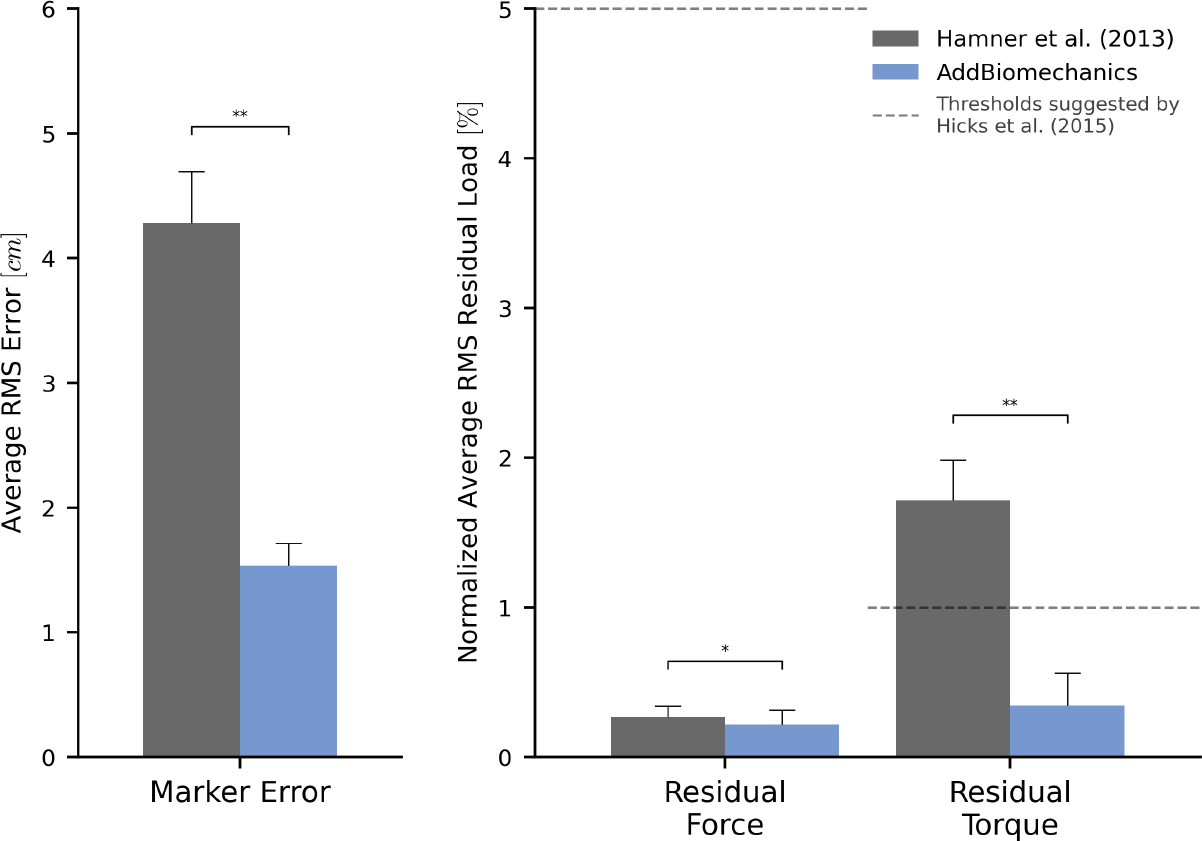
Human expert versus automated processing: running dataset. The root-mean-square marker errors (left) and residual forces and torques (right) from the original published study from Hamner et al. [62] (gray) compared to the results obtained using AddBiomechanics (blue). The results from Hamner et al. [62] were obtained using OpenSim’s scaling, inverse kinematics, and inverse dynamics tools, and residual loads were minimized using OpenSim’s Residual Reduction Algorithm (RRA). The residual forces are normalized to a percent of the peak ground reaction force, and the residual torques are normalized to a percent of the peak ground reaction force times the average center of mass height. The solid bars show the average per-trial RMS error, averaged over the 10 subjects in the evaluation. The error bars show the standard deviation of RMSE across the subjects. The dashed horizontal lines represent residual force and torque magnitude thresholds recommended by Hicks et al. [34]. Asterisks indicate statistical differences based on pairwise t-tests.

**Fig 4.**
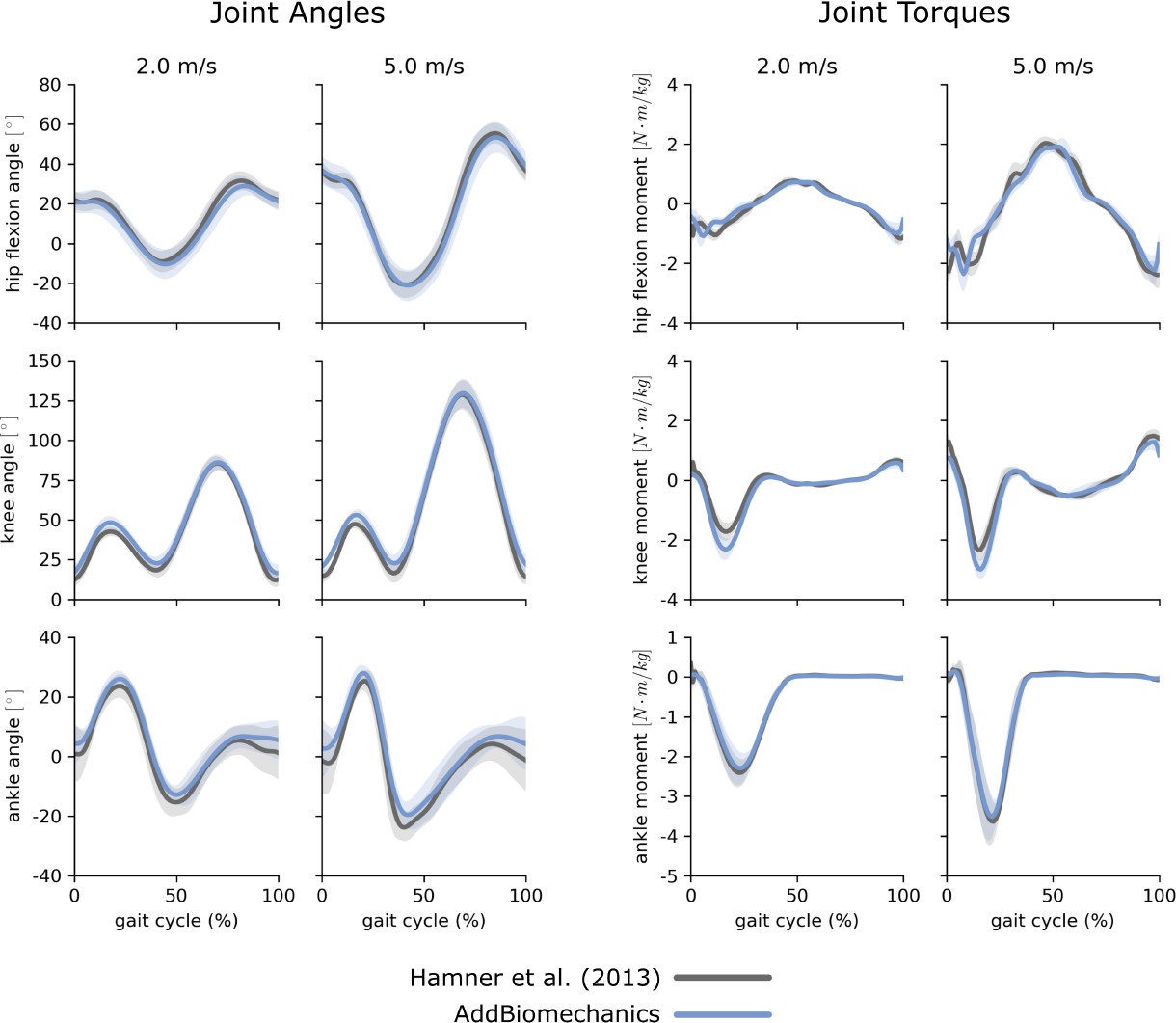
Running data: joint angles and torques. Joint angles (left) and joint torques (right) from the original published study from Hamner et al. [62] (gray) compared to the results obtained using AddBiomechanics (blue) for the 2.0 and 5.0 m s^-1^ running trials. The solid lines represent joint angles and torques averaged over the 10 subjects in the evaluation; the shaded bands represent the standard deviation across subjects.

The manual data processing by the expert in the original publication was labor intensive: each participant took several days for the expert to create a dynamically-consistent scaled model and compute joint angles and torques. Average computation time for a participant processed with AddBiomechanics was less than 30 minutes on a desktop machine, with 3-5 minutes spent on scaling and inverse kinematics, and the remainder on dynamic consistency.

### Human expert versus automatic processing: multi-activity dataset

AddBiomechanics produced similar marker errors (RMS: 1.6 cm, max: 3.9 cm) when processing the multi-activity dataset compared to manual processing by experts (RMS: 17 cm, max: 3.7 cm; Fig 5, left). The original study published by Uhlrich et al. [49] did not perform a residual reduction step before computing joint moments. However, AddBiomechanics automatically produced an inverse dynamics solution that met the recommendations of Hicks et al. [34] (Fig 5, right) and significantly reduced both residual forces and moments (p *<* 0.005, paired t-test). In addition, the lower-limb joint angle and joint torque trajectories from the automated approach were qualitatively similar to the trajectories from the original study (Fig 6).

**Fig 5.**
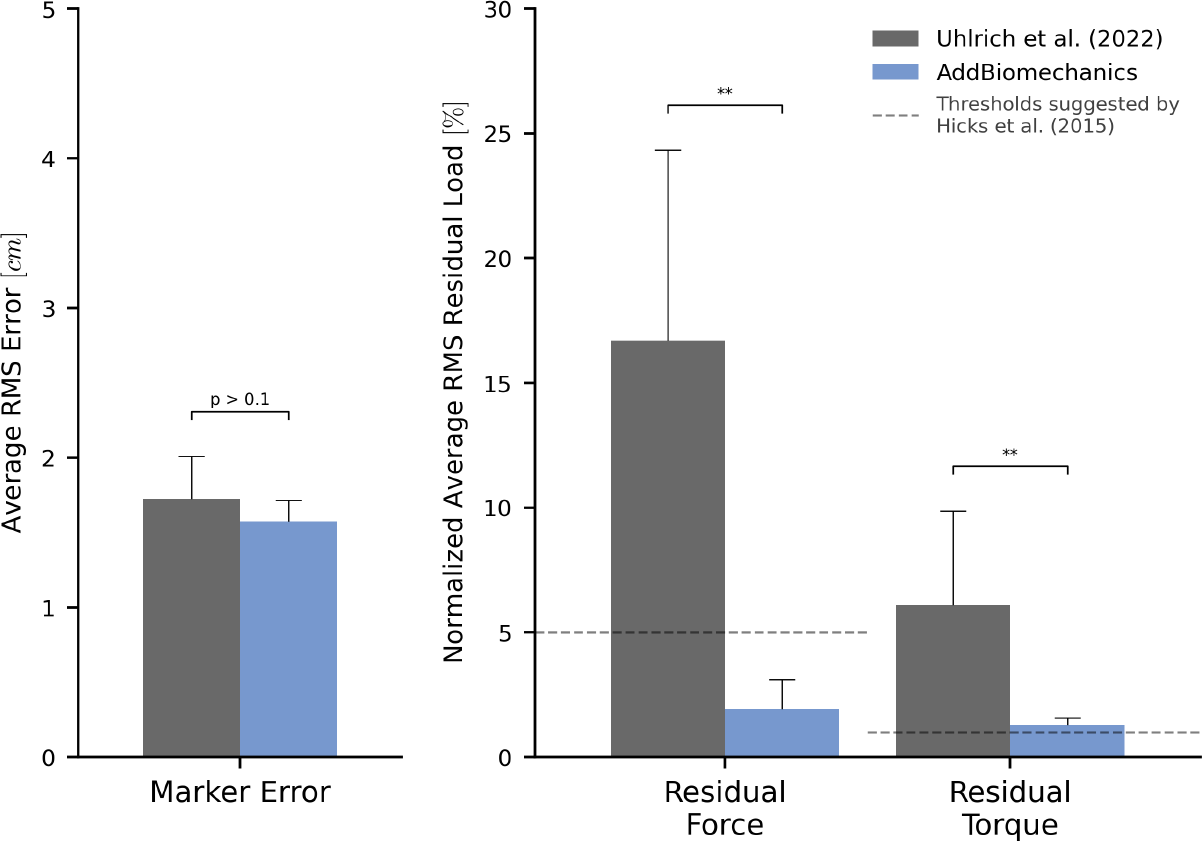
Human expert versus automatic processing: multi-activity dataset. The root-mean-square marker errors (left) and residual forces and torques (right) from the original published study from Uhlrich et al. [49] (gray) compared to the results obtained using AddBiomechanics (blue). The results from Uhlrich et al. [49] were obtained using OpenSim’s scaling, inverse kinematics, and inverse dynamics tools, but no residual reduction step was performed. The residual forces are normalized to a percent of the peak ground reaction force, and the residual torques are normalized to a percent of the peak ground reaction force times the average center of mass height. The solid bars show the average of per-trial RMS error, averaged over the 10 subjects in the evaluation. The error bars show the standard deviation of RMSE across the subjects. The dashed horizontal lines represent residual force and torque magnitude thresholds recommended by Hicks et al. [34]. Asterisks indicate statistical differences based on pairwise t-tests.

**Fig 6.**
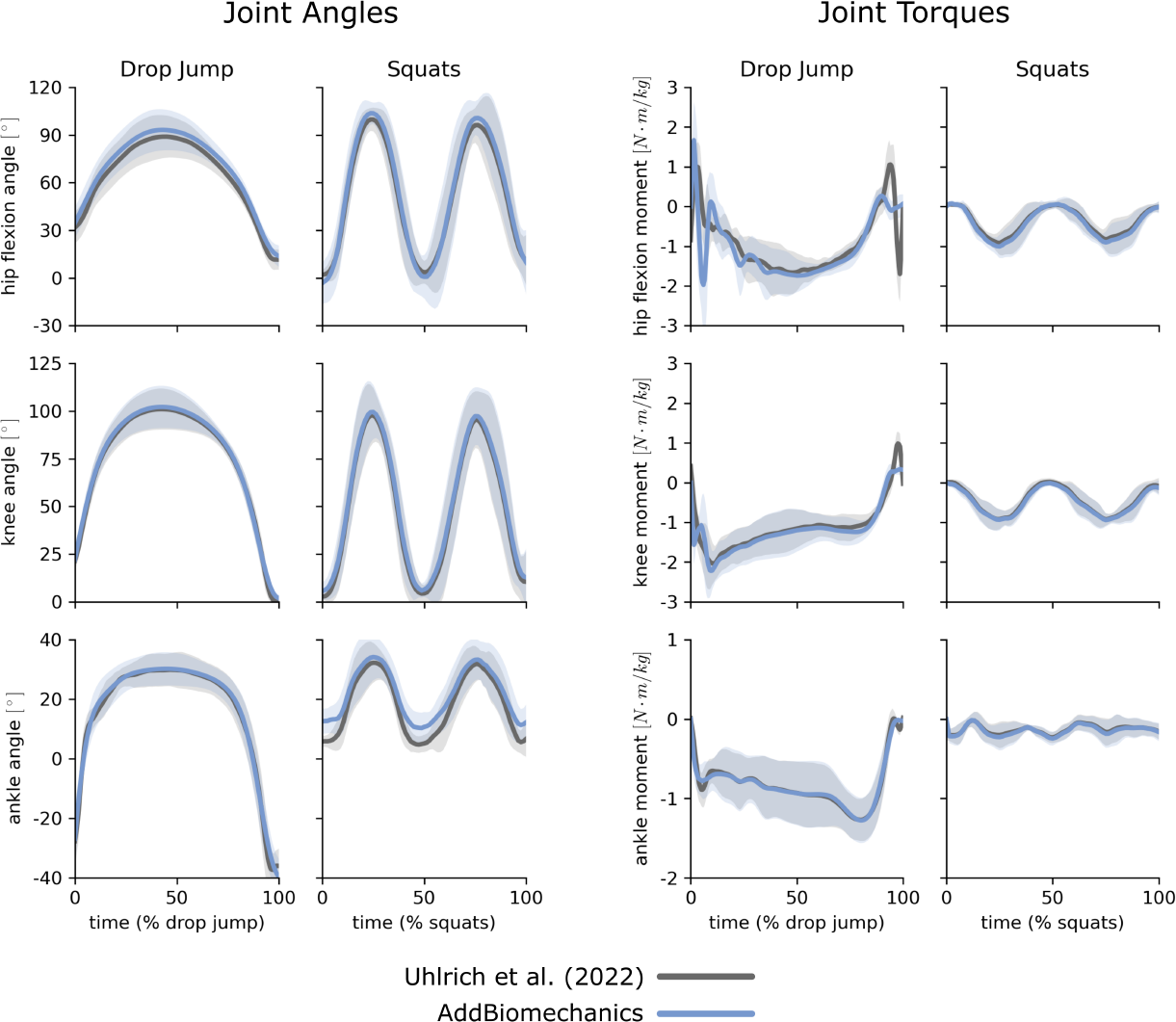
Multi-activty data: joint angles and torques. Joint angles (left) and joint torques (right) from the original published study from Uhlrich et al. [49] (gray) compared to the results obtained using AddBiomechanics (blue) for drop jump and squatting activities. The solid lines represent joint angles and torques averaged over the 10 subjects in the evaluation; the shaded bands represent the standard deviation across subjects.

Manual expert scaling for the multi-activity dataset was also labor intensive, taking roughly one working day per subject, not including additional time to perform inverse kinematics and inverse dynamics for each of the movement trials. AddBiomechanics required less than one hour on a desktop machine to automatically perform scaling, inverse kinematics, and inverse dynamics for each subject with no input from the user. Scaling and inverse kinematics was completed in under 10 minutes, with the remaining time being consumed by dynamics processing.

### Synthetic walking data results

We found that AddBiomechanics was able to recover the ground truth joint angles and joint torques from the synthetic walking marker data to an average of 1.6 deg RMSE and 0.15% body weight times height (computed over all joints in a trial together). The marker errors and residual loads achieved by AddBiomechanics for the synthetic data were small (0.63 cm and 0.01% normalized load, respectively; Table 1).

**Table 1.** Synthetic walking data results.

| Quantity | Average RMSE<sup>†</sup> | Units |
| --- | --- | --- |
| Joint Angles | 1.6 ± 0.3 | degrees |
| Joint Torques | 0.15 ± 0.01 | % BW × height |
| Markers | 0.63 ± 0.08 | centimeters |
| Residual Force | 0.01 ± 0.01 | % normalized force<sup>‡</sup> |
| Residual Torque | 0.01 ± 0.01 | % normalized torque<sup>‡</sup> |

<sup>†</sup> The average RMSE results are presented as mean ± standard deviation across subjects. <sup>‡</sup> The residual forces are normalized to a percent of the peak ground reaction force, and the residual torques are normalized to a percent of the peak ground reaction force times the average center of mass height.

## Discussion

Our bilevel optimization algorithm to find body segment scales, marker offsets, and joint angle and torque trajectories found dynamically-consistent trajectories for the multi-activity dataset while achieving marker reconstruction errors similar to the originally published expert-processed data. In addition, AddBiomechanics was able to automatically reproduce lower-limb joint angles and torques from the running dataset while achieving similar residual loads and significantly reducing marker error. Finally, AddBiomechanics reproduced the joint angles and torques from the synthetic walking dataset with high accuracy while achieving very low marker error and residual forces. The sequential approach we used to create initial guesses for solving the model scaling, inverse kinematics, and inverse dynamics optimizations problems made our method fast and robust, requiring no expert intervention.

In addition to being computationally efficient, our method improves upon previous automated model optimization methods. For comparison, the method in [24] assumed that all the body segment scalings were known to the algorithm and only attempted to find the marker offsets and the joint angles, and resulted in 1.21 degree joint angle RMSE. Our method must also recover segment scaling information from the data but achieves similar results: processing the synthetic walking data led to a joint angle RMSE of 1.6 degrees. The marker error results from our approach are also consistent with previous automated scaling approaches, which all outperform human experts when fitting a model to the same data [24, 43, 44, 64–66]. However, previous approaches required large amounts of compute time, were limited to one specific skeleton, or only addressed part of the body segment scaling and marker registration problem. In addition, our method found inverse dynamics solutions with normalized residual forces and torques similar to the results from the automated RRA optimization algorithm proposed by Sturdy et al. [25]. Our approach found scaling, inverse kinematics, and inverse dynamics solutions for multiple trials in less than 30 minutes, whereas the approach by Sturdy et al. [25] can take up to two hours to find dynamics for a single trial, and requires scaling be known in advance.

Our optimization approach has some limitations that should be considered when processing experimental movement data with AddBiomechanics. First, there is some fundamental ambiguity in reconstructing the full kinematic and anthropometric information (body segment scales, marker offset registrations, and body positions) from only marker location data. For example, the pelvis can be tilted slightly forward, with the markers at the front of the pelvis shifted upward, and if the angles of the hips and spine are appropriately adjusted then the markers will still closely match the target data. If this effect is observed in practice, AddBiomechanics users can leverage the fact that the optimizer will prioritize solutions that move the anatomical markers as little as possible, and adjust the marker starting locations on the bones to more closely match the experimental placement. Second, the optimizer applies a statistical prior to body segment scales to bring them more in-line with population statistics as represented by the ANSUR II anthropometric dataset [55]. If the optimizer can find a way to fit the marker data with a skeleton that is more likely to exist in the ANSUR II population (such as by tilting the pelvis forward 2 degrees), it will choose that one, even if the “true” underlying skeleton was slightly different. The data in ANSUR II is large and detailed, but was collected from active-duty military personnel, and so is not reflective of many patient populations. A broader anthropometric dataset could help address this limitation. Finally, AddBiomechanics may not always find an inverse dynamics solution with sufficiently low residual forces and torques due to inconsistencies between the marker and ground reaction force data that cannot be accounted for with a rigid body model.

By creating and sharing this tool, we aim to make quantitative biomechanics results more accessible, including to clinicians and researchers who do not possess the technical expertise or time traditionally required to achieve high-quality results. Our method goes from labeled marker trajectories to a scaled, registered, and physically-consistent musculoskeletal model and corresponding human motion in less than 30 minutes on a low-end server. We also provide a web version at AddBiomechanics.org which features a drag-and-drop interface to automatically process human movement data in the cloud. In exchange for sharing the resulting anonymized motion data with the scientific community under a creative commons license, we make AddBiomechanics freely available for researchers. As of this writing, over 300 researchers have used the prototype tool to process and share more than 14,000 motion files from almost 1,200 experimental subjects. We hope AddBiomechanics will increase the quality, consistency, and availability of biomechanical data analyses and lead to the creation of a large-scale public dataset of accurately modeled human motion biomechanics.

## Data availability statement

All data and code used for running experiments, model fitting, and our cloud application is available on a GitHub repository at https://github.com/keenon/AddBiomechanics and we have archived our code on Zenodo (DOI: 10.5281/zenodo.6981568). The data and code used to generate the results can be found at https://github.com/stanfordnmbl/addbiomechanics-paper.

## Acknowledgments

KW and MR received support from the National Science Foundation Graduate Research Fellowship Program (DGE-1656518; https://www.nsfgrfp.org/). NAB, JLH, SLD, and CKL received support from the Wu Tsai Human Performance Alliance at Stanford University and the Joe and Clara Tsai Foundation (https://humanperformancealliance.org/). MR, SHC, and CKL received support from Stanford Human-Centered Artificial Intelligence, Hoffman-Yee Research Grant program (Stanford-HAI-203112; https://hai.stanford.edu/). JS received support from the Stanford Bio-X SIGF Paul Berg Interdisciplinary Biomedical Graduate Fellowship (https://biox.stanford.edu/). JLH and SLD received support from the National Institutes of Health (P41-EB027060 and P2C-HD101913; https://www.nih.gov/). The funders had no role in study design, data collection and analysis, decision to publish, or preparation of the manuscript. Special thanks to Carmichael Ong, who lent us his OpenSim expertise and helped create several skeletons and markersets for AddBiomechanics, and Reed Gurchiek for early feedback on the tool.

## Supporting information

### S1 Appendix

#### Joint acceleration smoothing

Prior to finding inverse dynamics solutions with AddBiomechanics, we first perform a simple optimization to smooth the inverse kinematics solution we found by minimizing the jerk in the joint angle trajectories. This step is necessary to prevent large acceleration artifacts that appear as a result of small differences in joint angles between adjacent time steps from inverse kinematics. This optimization computes a new set of joint angles, 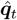, and includes a regularization term controlled by the weighting parameter, *σ*, which prevents large deviations from the original inverse kinematics solution, ***q***_*t*_.

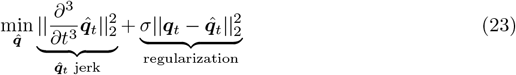

In our implementation, the joint jerks 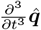 are computed using finite differences.

#### Constructing the 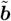 vector for angular dynamics fitting

The vector 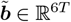 represents the current COM trajectory through space (first half, 3*T* entries), and the current root (e.g., pelvis) angular trajectory (second half, 3*T* entries), at the initial conditions ***ζ***.

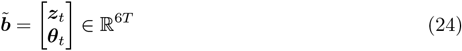

To compute ***z***_*t*_, note that we are holding the mass of the subject constant during this optimization step (*m*), so we can simply integrate the effects of known external forces (gravity, GRFs) on the known mass of the subject over time.

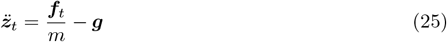

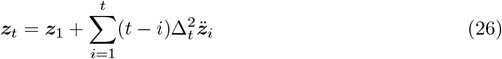

To compute ***θ***_*t*_, we linearly integrate the “residual free angular acceleration” over time. We define the “residual free angular acceleration” with joint state ***q***_*t*_, 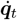, 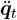 to be the necessary root angular acceleration 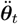 such that no angular residual force is present. By convention 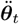 is always the first three entries of 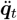, which we write 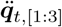.

We can compute 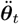 given ***q***_*t*_, 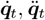 by first solving for joint torques ***τ***_*t*_ using inverse dynamics. Then, set the first three entries of ***τ***_*t*_ to 0, and solve forward dynamics using ***q***_*t*_, 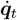, ***τ***_*t*_. The first three entries of the resulting 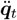 are the “residual free angular acceleration” 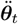.

Given the “residual free angular acceleration” 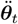, we can compute ***θ***_*t*_ by linearly integrating the “residual free angular acceleration” over time:

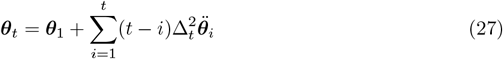

#### Accounting for force plate registration errors

The location of experimental force plates are often registered incorrectly in the motion capture volume, which leads to solutions with center of mass trajectories offset slightly from the experimental marker data. To align our solution with the observed experimental data, we shift both the location of the force plates and the marker trajectories by the offset between the center of mass trajectory we find and the experimentally estimated center of mass trajectory, which typically is only a fraction of a centimeter.

It is also common to have ground reaction force data recorded from force plates that are very slightly (e.g., less than 0.5 degrees) off of perfectly vertical. This makes finding a physically-consistent solution challenging, because if we assume that the force plates are perfectly vertical in our optimization problem, the total ground reaction force vector will be very slightly off of perfectly vertical in the ground reference frame. This can lead to a substantial horizontal acceleration bias on very long trajectories, since the horizontal ground reaction forces will not be exactly anti-parallel to gravity.

Since the force plate rotation errors are typically very small, we can preserve the linearity of our system by using a first-order Taylor expansion to approximate the rotation of measured forces by very small angles. In our implementation, we append a rotational correction term ***α***_*i*_ *∈* ℝ^3^ to ***ζ*** for each force plate *i*, where *n* is the number of force plates.

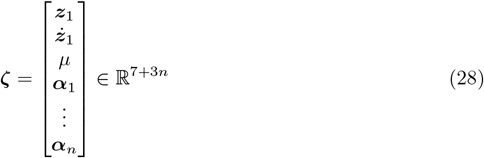

For each ***α***_*i*_, we append to ***A*** a 3*T ×* 3 columnar block:

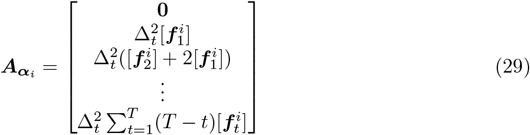

where 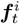 is the ground reaction force vector associated with force plate *i*. The [.] operator makes a skew-symmetric matrix out of a vector in ℝ^3^, so that ***a*** *×* ***b*** = [***a***]***b***.

Finally, we regularize to angles ***α***_*i*_ to discourage large force plate rotations. With this extension to our linear fitting problem, we can recover force plate rotations to a very small fraction of a degree.

#### Quantifying the impact of using the bilevel constraint in kinematics fitting

We ran an ablation study to quantify the impact of the formulating our kinematics fitting problem as a bilevel optimization problem. Here, we compare two conditions: optimizing a bilevel objective and optimizing a simple monolevel objective. The objective for the bilevel optimization problem (stated previously in Eq (2)) is as follows:

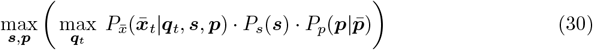

To convert the bilevel objective to a monolevel objective, we move joint angle optimization, max_***q****t*_, to the outer objective along with the body scale and marker registration optimization:

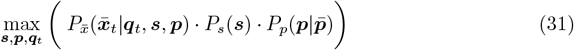

We find that using a bilevel approach leads to much faster convergence, and we can get lower marker RMSE at the cost of slightly lower anthropometric prior probability. The bilevel optimizer takes slightly more wall-clock time per iteration, but because it is able to reach high quality marker RMSE in many fewer iterations, it is able to save wall-clock time overall. We ran both optimization functions for several different fixed numbers of iterations on a single walking trial on a commodity server; these results are summarized in Table A1.

**Table A1.**
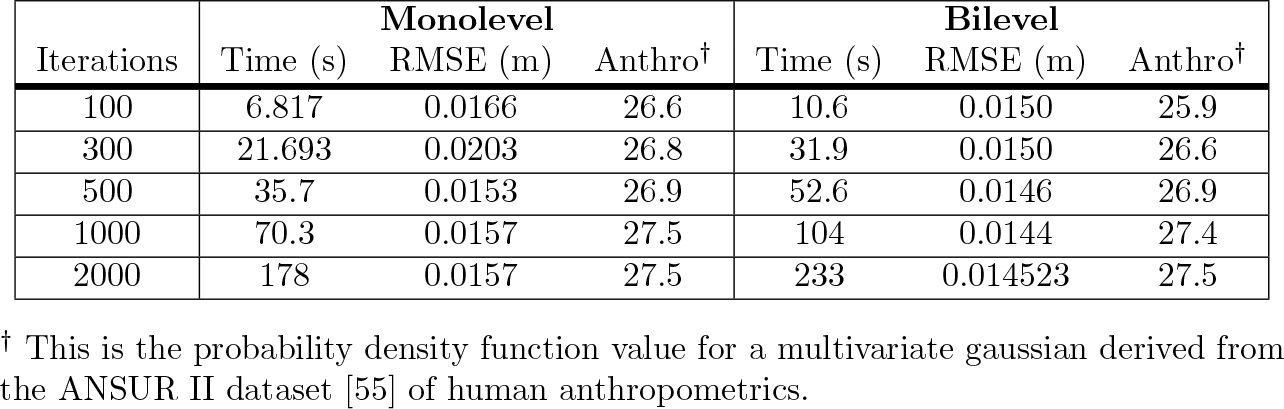
Monolevel versus bilevel optimization.

| Iterations | Monolevel |  |  | Bilevel |  |  |
| --- | --- | --- | --- | --- | --- | --- |
|  | Time (s) | RMSE (m) | Anthro<sup>†</sup> | Time (s) | RMSE (m) | Anthro<sup>†</sup> |
| 100 | 6.817 | 0.0166 | 26.6 | 10.6 | 0.0150 | 25.9 |
| 300 | 21.693 | 0.0203 | 26.8 | 31.9 | 0.0150 | 26.6 |
| 500 | 35.7 | 0.0153 | 26.9 | 52.6 | 0.0146 | 26.9 |
| 1000 | 70.3 | 0.0157 | 27.5 | 104 | 0.0144 | 27.4 |
| 2000 | 178 | 0.0157 | 27.5 | 233 | 0.014523 | 27.5 |

<sup>†</sup> This is the probability density function value for a multivariate gaussian derived from the ANSUR II dataset [55] of human anthropometrics.

